# Attila and Aetius on the roof: The succession phenomenon in the roof greening area of a primary school in Beijing during the lockdown period of the COVID-19

**DOI:** 10.1101/2023.11.07.565953

**Authors:** Ruoxi Wu, Jingqi Guan, Zihang Xia, Tianqing Zhang, Yi Jiang, Zhong Wang

## Abstract

“Succession” refers to the process of certain species replacing others over time. It is of great significance for understanding the characteristics and evolution of ecosystems. This article compares the changes in school rooftop green areas before and after the COVID-19 epidemic (2019, 2023), and studies the situation of wild grass invading roofs during the three years of the epidemic, as well as the response of original rooftop plants to the invasion. The results showed that: ① invasive weeds were concentrated in 2 families and 4 species, forming a population advantage in most of the invaded planting boxes; ② The succession has had a significant impact on the roof greening area; ③ Among the two cultivated plants (*Phedimus aizoon* (PA) and *Sedum sarmentosum* (SS)), SS was greatly affected, with the community almost disappearing, while PA was almost unaffected.

## 1. Introduction

Succession refers to the sequential development and changes in community composition and environment over time, as some species invade and others disappear in a biological community. Community succession is an important means of understanding the relationship between biodiversity and ecosystem function (Allan Raffard et al., 2018), and it is also a major tool for understanding community stability (Zheng Yuanrun, 2000). At present, research on succession phenomena has mainly focused on ecological restoration in ecologically degraded areas (such as the ecological restoration of grasslands in Inner Mongolia) (Wang Wei et al., 1996), as well as explanations for ecosystem changes and differences (such as differences in different water bodies in Canada) (Jeo S Schacht, 2023). Due to the environment of cities are always artificial, their green spaces are often artificial and fragmented, and there is relatively little research on them.

On the other hand, the arrival of the COVID-19 epidemic and the subsequent urban management and control provide a rare opportunity for the natural succession of cities. Although the epidemic has not completely passed, relevant research has already emerged. Such as, E. S. Diamant et al. studied bird behavior in the Los Angeles area and found that the Flight Initiation Distance of the black eyed sparrow (Dark eye Juncos) during the epidemic was significantly shorter, suggesting that due to the lockdown of the epidemic, birds have become less afraid of humans (E. S. Diamant et al., 2023). However, Congnan Sun et al. depicted a more dynamic and detailed picture: the activity of species heavily dependent on humans (such as the hooded crow) decreased significantly, while the activity of wild species adapted to urban life (such as the graceful prinia) increased, showing a complex succession pattern (Congnan Sun et al., 2023). To sum up, the COVID-19 epidemic may provide us with an excellent opportunity to observe the urban succession. However, throughout the literature, relevant research is still fragmented and lacks direct evidence of community succession.

Undoubtedly, the roof greening areas in large cities are a typical micro landscape ecosystem under human intervention, in which the animal, plant, and ecological environment foundations heavily rely on human intervention and support. This urgent need for artificial support is reflected in two aspects: ① Strictly speaking, even large rooftop green areas still belong to urban ecosystems, so they are inevitably different from the natural environment. “In terms of life support resources, these urban industrial environments are parasites in the biosphere” (E P. Odum, 2009, p366). Therefore, in the environmental context of large cities, artificially selected plants (which are obviously beneficial for improving the urban environment) often cannot adapt well to the microclimate of rooftops, especially in mega cities like Beijing located in climate transition zones (warm temperate continental monsoon → cold temperate continental monsoon). Previous studies have shown that only 2 out of 10 species of Sedum plants in the roof area have survived for 8 years (Hui Zhang et al., 2021) ② Artificial support can have a significant impact on competition among species populations. For roof greening areas, human activities such as weeding can significantly hinder the invasion of native wild plants on rooftop plants (Hui Zhang et al., 2021). Existing studies also support this point, such as Huang Defeng’s study on eutrophication in constructed wetlands, which shows that reasonable design and plant selection can effectively hinder eutrophication (Huang Defeng, 2007). It can be said that this is an absolutely artificial environment.

However, the occurrence of COVID-19 eliminated this fundamental factor. Taking the rooftop green area where this article was tested as an example, due to the three years of epidemic lockdown, the rooftop area was completely exposed to the natural environment of Beijing, not only without human care, but also facing competition from local wild plants. So, what happened to the roof area at this time? Will it completely degenerate into a pure wild environment? Observing and studying changes in the rooftop area at this time not only helps pupils of the school to understand changes in the ecological environment (such as population competition) and deepen their ecological awareness, but also provides a rare opportunity for monitoring and researching the control variables of artificial landscape environmental degradation. Furthermore, if a certain equilibrium and stability can be observed in the population competition between artificially selected plants and local wild plants, it may provide indirect evidence for the environmental improvement work in degraded ecological areas in China (such as the northwest sandstorm and arid areas that are difficult to sustain artificial care) (Feng Yi et al., 2009).

## 2. Species of the invasive plants and their diversity

### 2.1 Experimental area

The experimental area of this study is the rooftop area of a primary school in Dongcheng District, Beijing. Dongcheng District is the core city jurisdiction of Beijing, with a total area of 41.84km^2^, a permanent population of 704k (2022), and a GDP of about 47 billion dollars (2022). The government of Beijing, world cultural heritage sites such as the Forbidden City and Temple of Heaven Park, as well as world-renowned commercial pedestrian streets such as Wangfujing Pedestrian Street, are all located in this area. So, this area can represent the basic ecological characteristics of Beijing urban area.

During the 8-year period from 2013 to 2021, a research team from the Beijing Academy of Agriculture and Forestry Sciences conducted a long-term tracking experiment on the adaptability of roof plants in the rooftop green area of the primary school of this article (Hui Zhang et al., 2021). The experimental area of this article is located in 24 planting boxes within the green area. A comparative experiment was conducted in 2019 (the year before the outbreak of the epidemic) on the planting and stem segment sowing of two plants (Yi Jiang et al., 2022). The comparability of data before and after the epidemic is strong (see Figure 1 for the layout of the experimental area).

**Figure 1:**
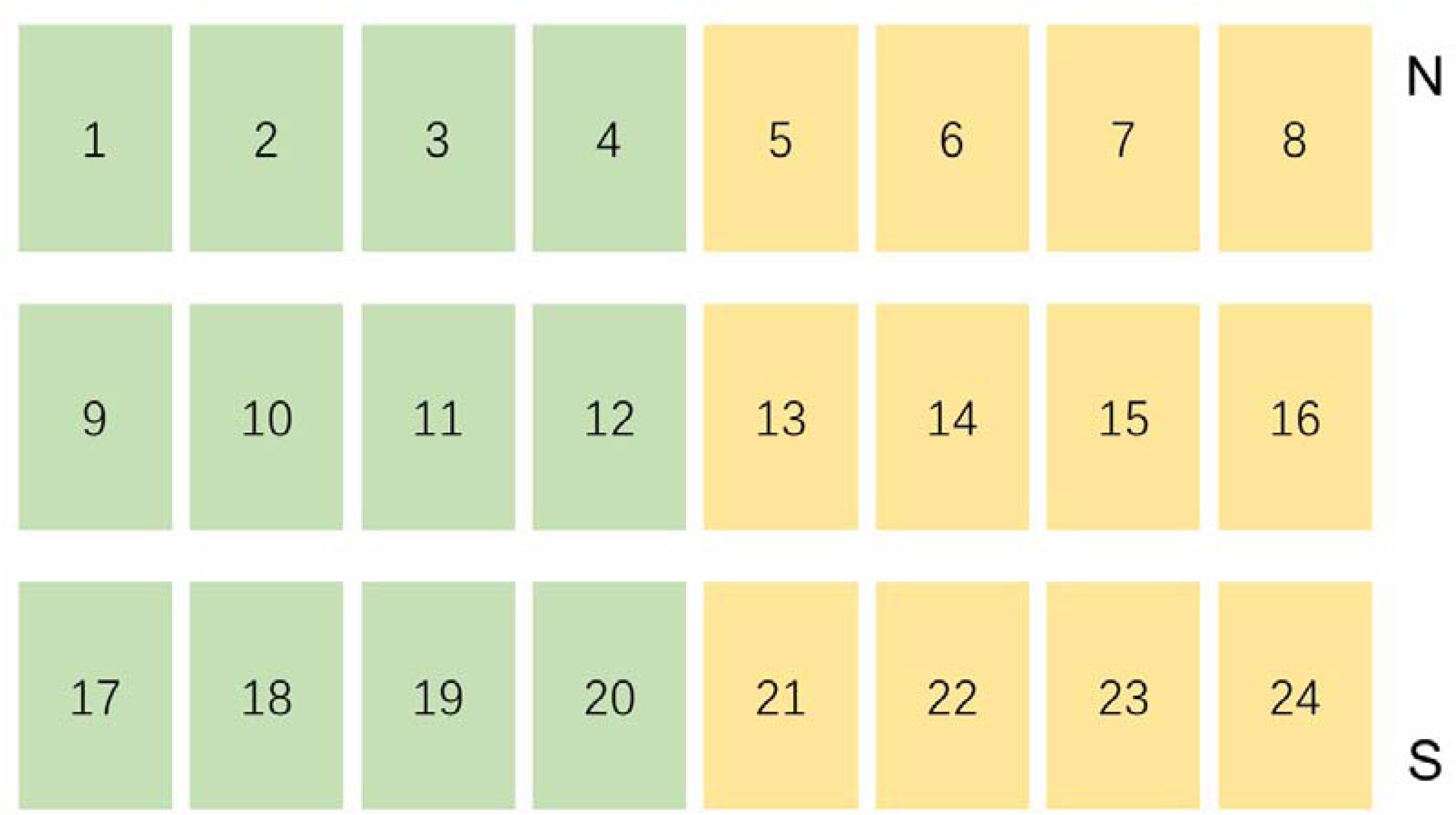
Overall layout of the experimental area (green boxes planted PA and with boxes planted SS)

### 2.2 Species of the invasive plants

Figure 1 shows the general layout of 24 planting boxes, and Figure 2 is a photo of the overall situation. From Figure 2, it can be seen that half of the planting boxes have suffered from severe invasion (mainly concentrated in the area of SS).

**Figure 2:**
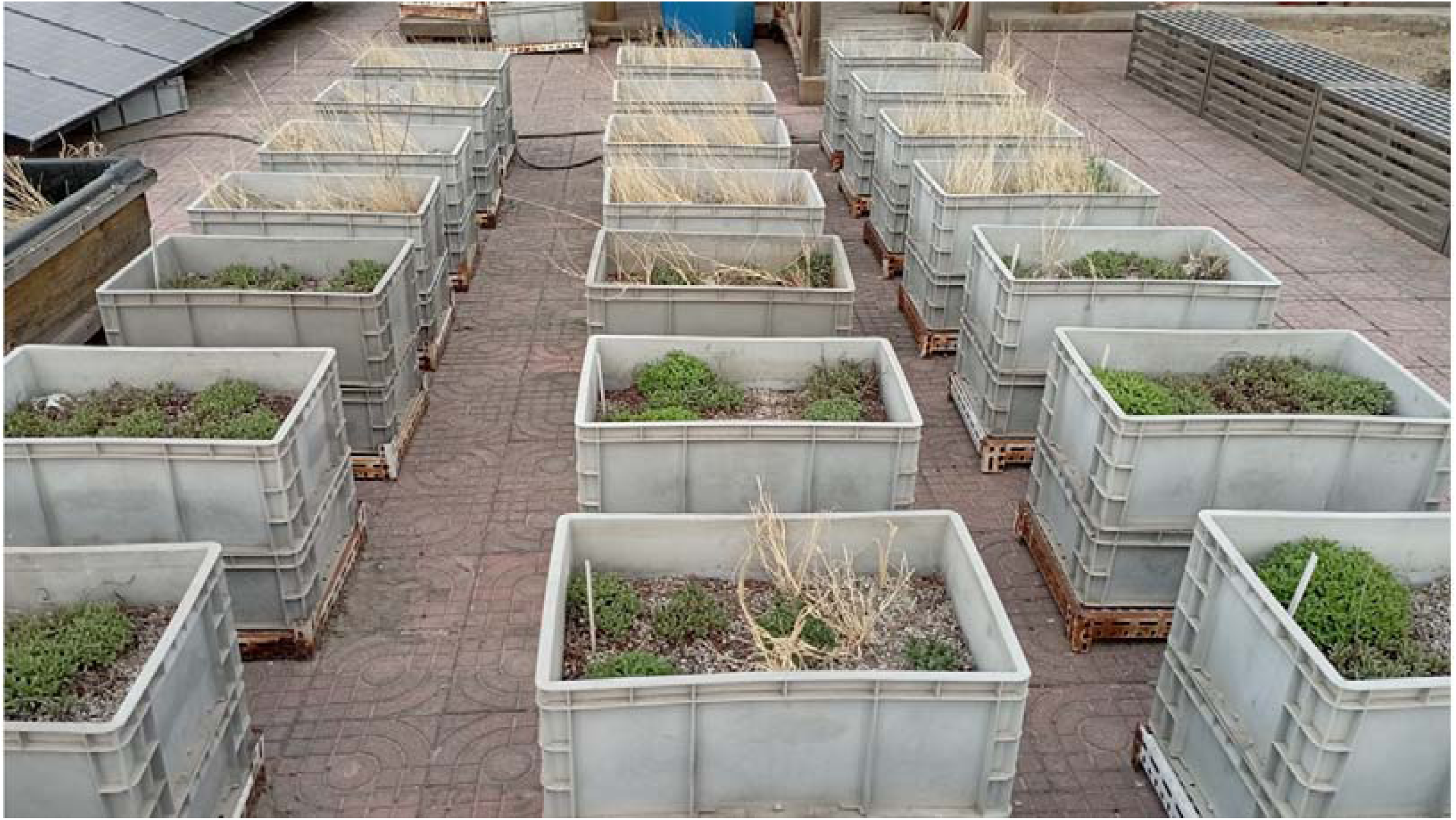
Site photos of the experimental area, showing that the SS area (in the distance) has been infected by a large number of wild grasses

We collected and produced specimens of invasive plants, and identified them using AI software combined with plant atlases. The artificial intelligence software chooses “Baidu Identifying” because relevant research shows that the software has a high accuracy rate among the common 8 software types in China (Zhanhui Xu et al., 2020) and is easy to use.

After morphological identification, the invasive plants are classified into the following four types:

i. *Avena fatua Linn. var. fatua* (AFL), an annual herbaceous plant of the Gramineae family, resembles oats and is one of the malignant weeds in farmland (Flora of China, wild oats);
ii. *Setaria viridis (L.) P. Beauv* (SVP), The annual herbaceous plant of the Gramineae family, Setaria, is widely distributed in temperate, warm temperate, and tropical regions of the world (Flora of China, Setaria);
iii. *Artemisia capillaris Thunb* (ACT). is a perennial semi shrubby herbaceous plant of the Artemisia genus in the composite family, with a strong fragrance. It is produced in most parts of China, mainly in Shanxi, Shaanxi, Anhui and other places (Flora of China, Artemisia annua);
iv. *Digitaria sanguinalis (L.) Scop* (DSS), The annual herbaceous plant of the Gramineae family, DSS, is widely distributed in temperate and tropical regions (Flora of China, Matang).

### 2.3 Diversity level of invasive plants

Due to the intermittent epidemic lockdown in schools from 2020 to 2022, and the fact that the rooftop green area is also under lockdown, so this article describes the outcome comparison (i.e., 2019 and 2023).

As mentioned above, 12 out of 24 planting boxes have suffered severe invasions. But as shown in Figure 2, the vast majority of planting boxes are occupied by a single invasive plant, which is in an absolute dominant position in the planting box. Only one planting box (23 #) has multiple plants, and we have calculated the area of each type of plant community in the planting box (Figure 3), as shown in the table below (see 3.1 of this article for the calculation method):

**Table 1.** Area of Each Plant in Box No. 23.

| ACT(P<sub>1</sub>) | DSS(P<sub>2</sub>) | AFL(P<sub>3</sub>) |
| --- | --- | --- |
| 0.145203 | 0.04 | 0.689145 |

**Figure 3.**
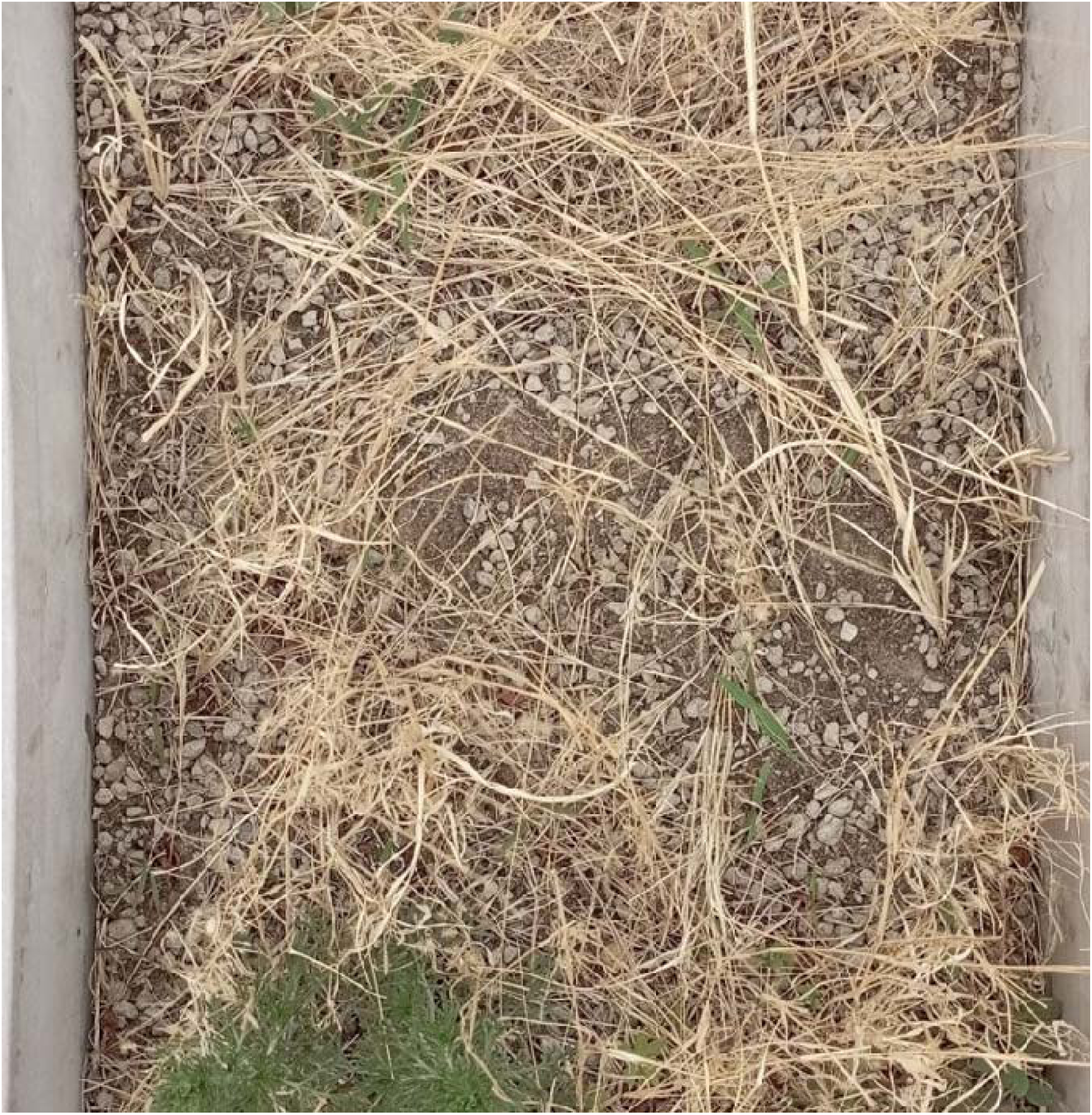
Box No. 23, where the ACT in the lower left corner and the DSS in the lower right corner can be clearly seen

Finally, the Shannon Wiener Index of the box is approximately

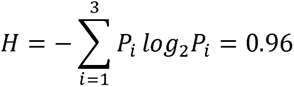

Note: Due to the large number of plants in the box, area is used instead of the number of plants.

### 2.4 Discussion

In this section, we investigated the main succession plants in the invasion experimental area, which are AFL, SVP, ACT and DSS. These four plant species are common in northern China, many of which are typical field weeds (Zhao Xinghua, 2017). It is worth noting that the types of wild grass collected this time are significantly less than those commonly found in the Beijing area. According to Zhao Xinghua’s survey of corn field grasses in the Huairou area of Beijing, the most common weeds were identified in 6 families and 9 species (Zhao Xinghua, 2017), while Lv Shihai’s survey of alfalfa fields in the Beijing area found a total of 16 families and 109 species of weeds (Lv Shihai, 2001). In addition to observation errors, there are two possible reasons why only four types of weeds were found in this article: firstly, the experimental area in this article is located on the roof, which leads to the need for weed propagation to rely on wind or bird drive. Therefore, those with higher seed weight are clearly not conducive to invasion; At the same time, this type of weed must adapt to the shallow soil depth of the roof, while weeds with well-developed roots such as *Arctium lappa* are clearly not suitable for the roof area. Secondly, this article mainly focuses on the observation of 2023, which means that there may have been several rounds of succession in the rooftop area between 2019 and 2023, and many weeds may have been eliminated. From Figure 2, it can also be clearly seen that AFL and SVP grass have a significant advantage in sunlight competition due to their high plants, while plants such as purslane may wither and rot if they cannot see sunlight for more than two days (Zhao Xinghua, 2017).

## 3. The overall pressure of invasion on the original ecology

Undoubtedly, this invasion has brought competitive pressure to the original artificial ecosystem, but how much is this pressure? Is it statistically significant? As mentioned earlier, due to the lockdown policy from 2020 to 2022, we can only observe the experimental area during the reopening period of 2023. Therefore, we are comparing the outcomes of the same region (pre pandemic vs. post pandemic).

### 3.1 Ideas and Methods

We compared the growth outcomes of two artificially planted plants in 24 planting boxes in 2019 and 2023.

Selected plant projection coverage as the main indicator. According to the methods of Ji Zengbao and Alexander Stewart, we used a method of taking photos of each planting box and quantifying the digital images to calculate the coverage of all planting boxes (Ji Zengbao, 2017) (Alexander Stewart, 2007). The specific method is to first took a vertical overhead photo of each planting box (equipped with a Redmi Note 11 phone rear camera with 50 million pixels), and then used the “Color Range” function in the “Selection” menu of Photoshop 2018 software to adjust its parameters to 120, so as to peeled off the vegetation and soil background color in the photo. Finally, used the “Histogram” function to calculate the ratio of the plant’s pixel count to the total pixel count in the image. Figure 4 shows the comparison before and after soil stripping in one of the planting boxes, and it can be seen that the effect is very obvious and the accuracy is high.

**Figure 4.**
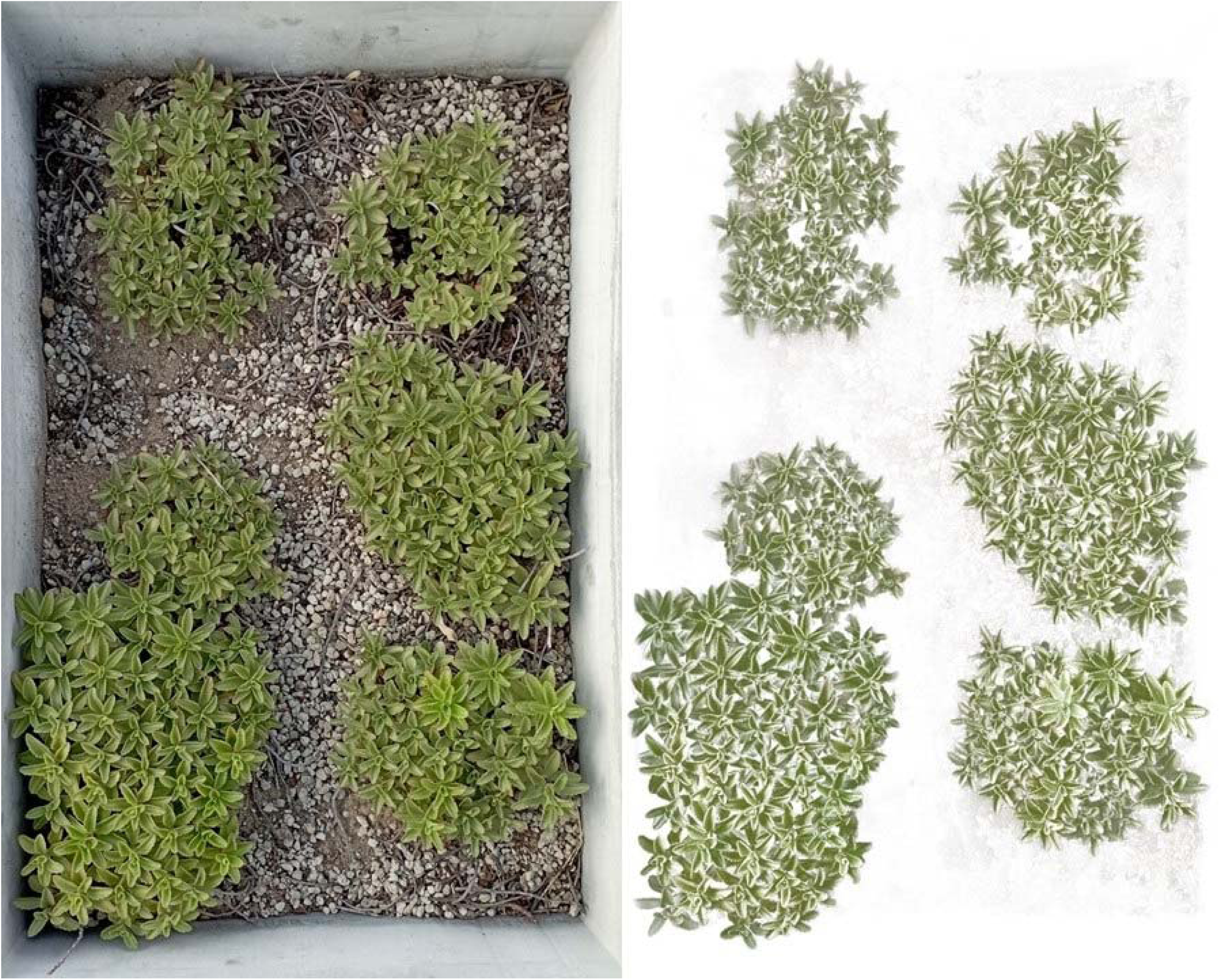
Comparison of One Planting Box Before and After Stripping the Soil Layer with Photoshop Software

After obtaining the coverage ratio of each image, we compared it with the control group data. The control group data was the same two species plants in the same region by Yi Jiang et al. in 2019 (Yi Jiang et al., 2023). The comparison method was independent sample T-Test (T-Test), and the statistical software was SPSS 24.0.

### 3.2 Results and Analysis

#### 3.2.1 Results

We obtained the projection coverage values for all 24 boxes in 2019 and 2023 using the above method, as shown in the table below

As shown in the table below, the difference is extremely significant (p<0.001), indicating a significant difference before and after the occurrence of succession, it means that the succession may have caused significant survival pressure on the experimental area.

#### 3.2.2 Excluding the data of SS

But if carefully observe the data of all projection coverage values (Table 2), it is not difficult to find that this pressure is likely mainly from SS, as SS has almost completely disappeared. Is this the case? We excluded the data of SS and used the T-Test to separately compare the data of PA in 2019 and 2023. The results confirmed our hypothesis, p=0.261, and the results were not significant, which cannot indicate that SS was severely affected by the invasion.

**Table 2:** the data of all projection coverage values.

| **boxs** | **2019** | **2023** |
| --- | --- | --- |
| 1 | 0.346 | 0.305 |
| 2 | 0.380 | 0.369 |
| 3 | 0.469 | 0.626 |
| 4 | 0.561 | 0.636 |
| 5 | 0.635 | 0.000 |
| 6 | 0.652 | 0.000 |
| 7 | 0.770 | 0.000 |
| 8 | 0.788 | 0.000 |
| 9 | 0.577 | 0.448 |
| 10 | 0.462 | 0.319 |
| 11 | 0.472 | 0.490 |
| 12 | 0.560 | 0.290 |
| 13 | 0.514 | 0.000 |
| 14 | 0.446 | 0.000 |
| 15 | 0.545 | 0.000 |
| 16 | 0.778 | 0.000 |
| 17 | 0.562 | 0.290 |
| 18 | 0.385 | 0.295 |
| 19 | 0.441 | 0.603 |
| 20 | 0.521 | 0.433 |
| 21 | 0.697 | 0.000 |
| 22 | 0.505 | 0.000 |
| 23 | 0.719 | 0.000 |
| 24 | 0.713 | 0.000 |

Note: The data is the coverage area of all communities in each planting box
divided by the total area of the planting box

**Table 3:** the difference of 2019 and 2023.

| Year | Average | *p* |
| --- | --- | --- |
| 2019 | 0.563 | < 0.001 |
| 2023 | 0.213 |  |

Nevertheless, we still found some signs of succession from the two years’ PA data: firstly, although the p-value is not significant, it is also relatively small, indicating the possibility of succession influence; Secondly, the coverage data for 2023 has significantly shrunk compared to 2019. The following figure (Figure 5) is a scatter plot obtained by dividing the 2019 data by the 2023 data. It can be seen that the majority of the data is distributed in areas>1, with an average value of approximately 1.21. This result shows that the average coverage in 2019 is higher than in 2023 (although the p-value is not significant).

**Figure 5:**
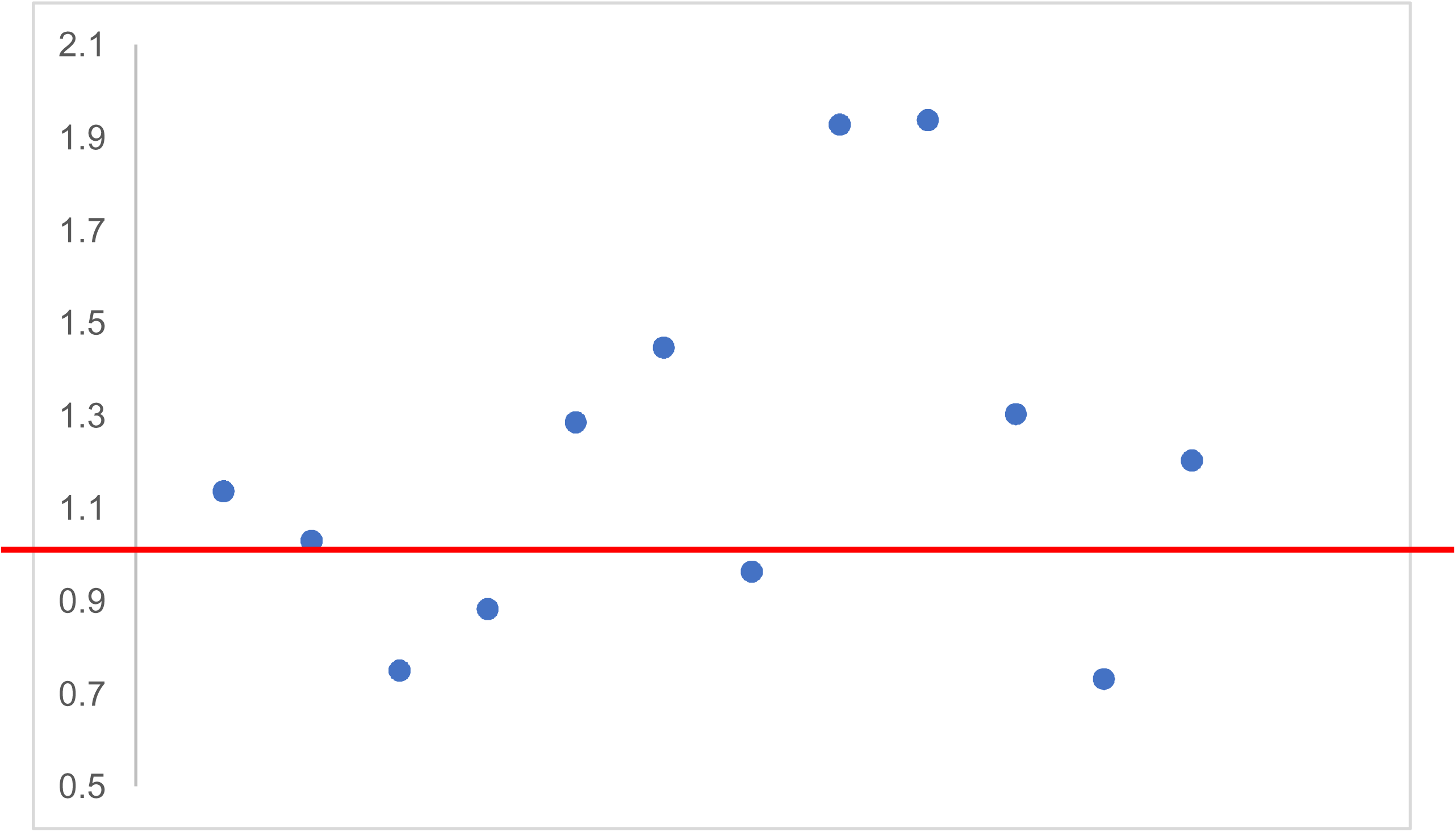
the ratio of SS coverage (2019/2023)

This result seems to indicate that PA may also be quietly changing, but the reason for this change is likely not from the invaders. To this end, we conducted chi-square tests on the data from 2019 and 2023 to test whether the coverage data of each planting box in the two years were closer to their average values (H_0_ was uniform distribution).

As shown in Table 4, the p-values for both years are greater than 0.05, indicating that they are relatively close to the average values of their respective groups. But it can be seen that the data for 2023 is closer to the average, in other words, the coverage data for each box in 2023 can be almost seen as a uniform distribution. It should be noted that before 2019, different densities of stem segments were sown in each box of this experimental area, which means that the original density of plants in each box was different (as required by the experiment). The comparison results seem to reflect a phenomenon: although PA was not affected by too many foreign invasions, they seem to be quietly trying to approach the average, that is, the box that were originally too dense became sparse, and what was originally sparse gradually became dense. So, isn’t this also a form of succession?

**Table 4:** The degree of deviation from the averages value of each year.

| Year |  |  | *p* |
| --- | --- | --- | --- |
| 2019 | 0.333 | 1 | 0.564 |
| 2023 | 0.000 | 1 | 1.000 |

### 3.3 Discussion

In this section, we compared the differences between before and after succession. The results showed that the difference between the two was extremely significant, and most of this difference was caused by SS. Although PA appears to have resisted the invasion of weeds, it also seems to be changing itself, with a trend towards returning to the average (although evidence is insufficient).

There is no doubt that invasion causes damage and change to existing ecosystems, which can be well explained by C. Darwin’s theory of natural selection and supported by some evidence. For example, Lohbeck M. et al. found in their study of the secondary succession phenomenon of tropical forests in Chiapas, Mexico that as succession progresses, especially with the continuous increase of biomass, resources become increasingly limited, Resulting in significant differences between secondary dominant species and primary dominant species, especially in terms of feature similarity (Lohbeck M. et al., 2013). For example, MacLane et al. conducted research on the succession of floodplains in Illinois, USA, and found that although the impact of herbaceous plants is not significant, woody plants are likely to have a lasting impact on local ecology due to the opportunities provided by flooding for species invasion (McLane CR et al., 2012).

## 4. Response of two original plants to invasion

As mentioned earlier, PA and SS showed completely different reactions, with SS almost completely annihilated, while PA still looked lush. So, is there an obvious difference in their reactions?

### 4.1 Differences between the two

We used coverage data from 2023 and divided them into two groups: “PA” and “SS”, and used T-Test to test the differences between the two.

It should be said that the results do not actually require the use of statistical software, and the magnitude of their differences can almost be determined with the naked eye. The use of statistical methods only makes the evidence more rigorous: p<0.001, the response of the two to invasion can be said to be vastly different.

### 4.2 Excluded A Competitive Interpretation

Although the difference between the two is surprisingly significant, there is a clear competitive interpretation here: the growth speed of PA is significantly faster than that of SS. The following figure (Figure 6) shows the comparison of the average coverage doubling months of two plants from the control group from 2013 to 2017 (Hui Zhang et al., 2021), where the histogram label data represents a doubling of coverage in the X month of the year compared to the beginning of the year data (i.e., the month which coverages were twice that of the beginning of the year). From the figure, it can be seen that the doubling-month of PA basically around April each year, while the value of SS was always after May. That is to say, can the huge difference between the two of above (4.1) was due to the slower growth of SS, not the pressure of succession? Obviously, this is a reasonable interpretation.

**Figure 6:**
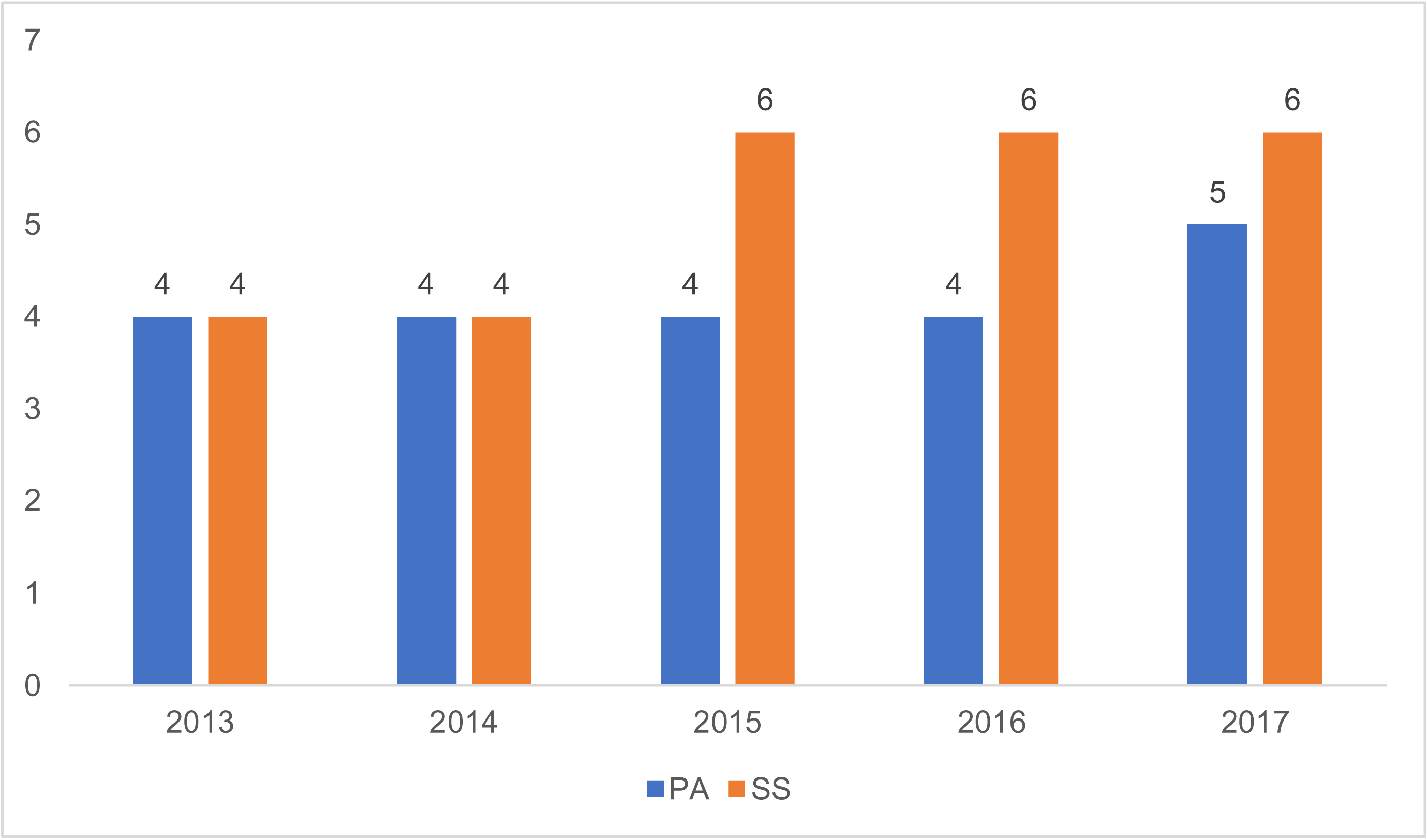
the doubling-month of the two species of original plants (2013-2017)

What truly overturned this idea was an accident: in mid-May, a school teacher unknowingly led some students to uproot the vast majority of visible weeds in the experimental area due to detest of disorderly weeds (Figure 7). This behavior objectively disrupted the experiment, but also unintentionally overturned this competitive interpretation: if the SS’s boxes were full of weed due to its slow growth, then now that all competitors have been removed, its growth process should be more easily observed. But the fact is obviously not like this. Until the end of the experiment in mid-July, only a few communities of SS showed their heads, and because the diameter of these barely surviving communities was less than 1cm, we did not record their coverage data.

**Figure 7:**
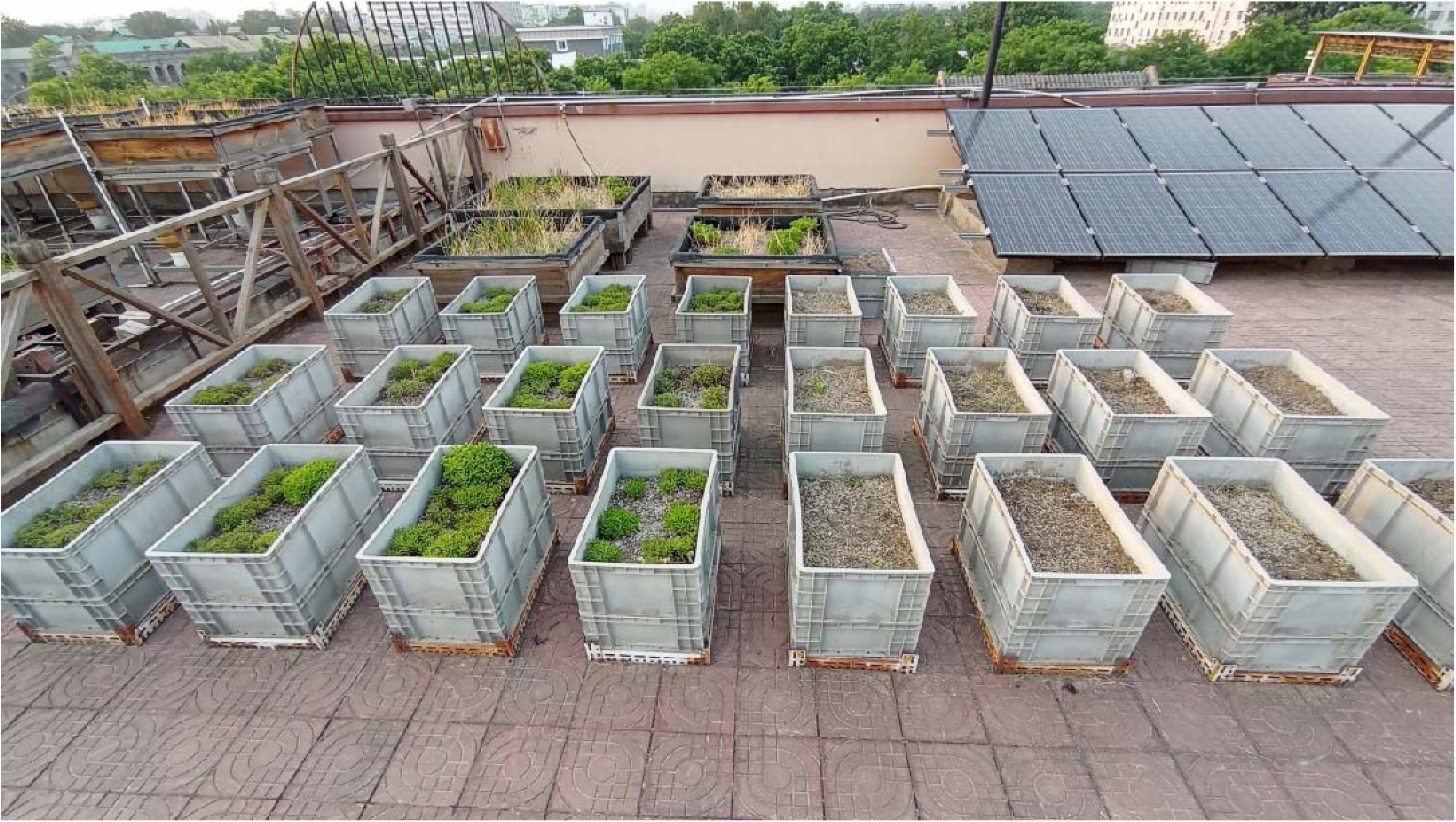
After the accidental removal of weeds in the experimental area. It is evident that there is no large-scale community of SS in the right 12 boxes(original SS area). Instead, there was a large area of bare soil, forming a sharp contrast with the PA area on the left (photographed on July 7, 2023)

### 4.3 Discussion

Although the relevant research is not rich, some existing evidence has shown that there are indeed some differences in the survival ability of the two. As pointed out in the control group literature of this article, during the eight year period from 2013 to 2020, PA survived, while the experimental SS all died (Figure 8, Hui Zhang et al., 2021). And, Yixin Song et al. conducted a 40-day drought resistance experiment on six species of Sedumaceae plants, and found that during severe water shortages of 30-40 days, PA’s survivability always ranked first or second, while SS was in the worst performing group (Yixin Song., 2023). The article further points out that the cold resistance of plants is closely related to their morphological characteristics (such as total leaf area and leaf shape), but not to their place of origin, that is, even plants from less arid areas are fully capable of adapting to arid environments (Yixin Song., 2023).

**Figure 8:**
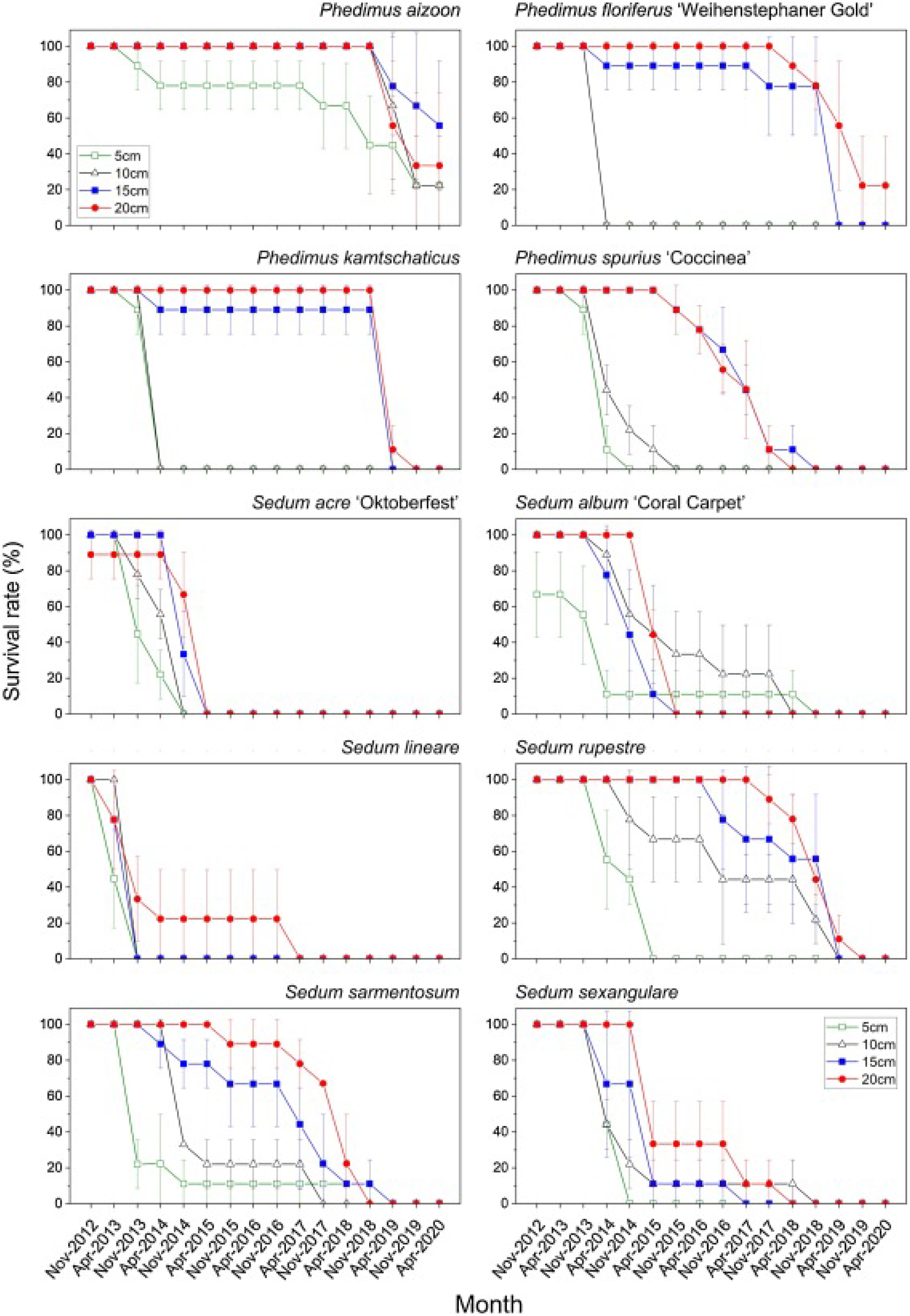
shows the survival rate of plants planted in the rooftop green area of the school by Hui Zhang et al., with the vertical axis indicator of the line chart showing the survival rate. From it, it can be seen that PA (upper left corner) still survived until 2020, while the data on SS (lower left corner) returned to zero, indicating that all of them had died.

## 5. General Discussion

This study observed and reported the succession of the roof green area of a primary school in Beijing during the three-year period of closure due to the lockdown of the COVID-19. In the first part of the study, we examined the main invasive weeds and found that they are mainly concentrated in 2 families and 4 species, and have formed absolute advantages in most of the invasive planting boxes. This was likely to indicate that the succession has undergone several rounds and was gradually stabilizing. In the second part, we compared the coverage data of 2019 and 2023 to examine the impact of succession on the originally planted plants, and found that the impact was significant. Further data analysis found that competition pressure mainly comes from the SS area (where the SS was almost completely destruction), while the PA area, although not significantly affected, was also undergoing succession, with the community density gradually tending towards the average. In the third part, we compared the response to succession between the originally plants. Although the data in the second part already indicates a significant difference in the response to competitive pressure between PA and SS, we further excluded a competitive interpretation in this part, that is, the difference shown in Figure 2 is not caused by the slow development of SS.

### 5.1 The huge impact of the COVID-19

The purpose of this paper is to illustrate that COVID-19 is a good window to observe the succession of ecosystems (especially urban green areas). The reason why this is an excellent period is that the COVID-19 epidemic provides a rare and unrepeatable external change force for succession.

The changes brought to the ecology during COVID-19 and the external pressure of succession on the artificial environment have already been discussed above, and will not be further discussed here. It is worth noting that this article is an observation and comparison of a micro ecological area with an area of less than 100 square meters and a time span of less than 3 years. This is likely to indicate that succession is also possible to develop and stabilize within such a short and narrow spatiotemporal region. This is not commonly seen in previous studies, and the area for succession research is rarely less than the entire roof of a building, and the observation period less than 5 years is also unusual. Therefore, the conclusion of this article may prompt a new understanding of ecology dynamics. Of course, these require further evidence to support them.

### 5.2 Remaining competitive explanations and shortcomings

There are still some competitive explanations in this article that cannot be answered through the above evidence:

Firstly, the biggest issue in this article is causality. As the second part of this article compares the changes in plant coverage in the same region between 2019 and 2023, we can be confident that the data can reflect the true situation of succession in these three years. But the question is, are the results of these successions (especially the complete annihilation of the SS) caused by weeds? Or is it due to some unknown reason that led to the large-scale death of the SS, allowing weeds to “take advantage of the situation”? Indeed, there is a possibility. According to the observation of the school’s teachers, crows living around the school occasionally peck at these two experimental plants, and according to their experience, crows seem to prefer pecking at SS, which is likely to be the reason for the downfall of SS. But this is just the experience of the teachers and there is no data support. Unfortunately, due to epidemic control, we are unable to enter the experimental area for observation between 2020 and 2023, so we lack relevant evidence to deny this possibility.

Secondly, weather is also a significant interfering factor. June this year was the hottest month on meteorological records, with major climate indicators including global air and sea surface temperature exceeding previous records (Katharine Sanderson, 2023). Francis, an expert at the Woodville Climate Research Center in the United States, called July 3rd the “hottest week in 100000 years” (Chinese meteorological enthusiast, 2023). And this will definitely reduce the reliability of the data and the persuasiveness of the conclusion in this article. As shown in Figure 9, a photo of one of the boxes on July 10th clearly shows signs of drying and withering. Due to the severe distortion of data caused by withering, all collected data in this article are due to the end of June. Therefore, this inevitably leads to another issue in this article: the data collection time in this study was significantly shorter than that of the control group by more than a month, and the persuasiveness of the conclusions in this article has been weakened to a certain extent.

**Figure 9:**
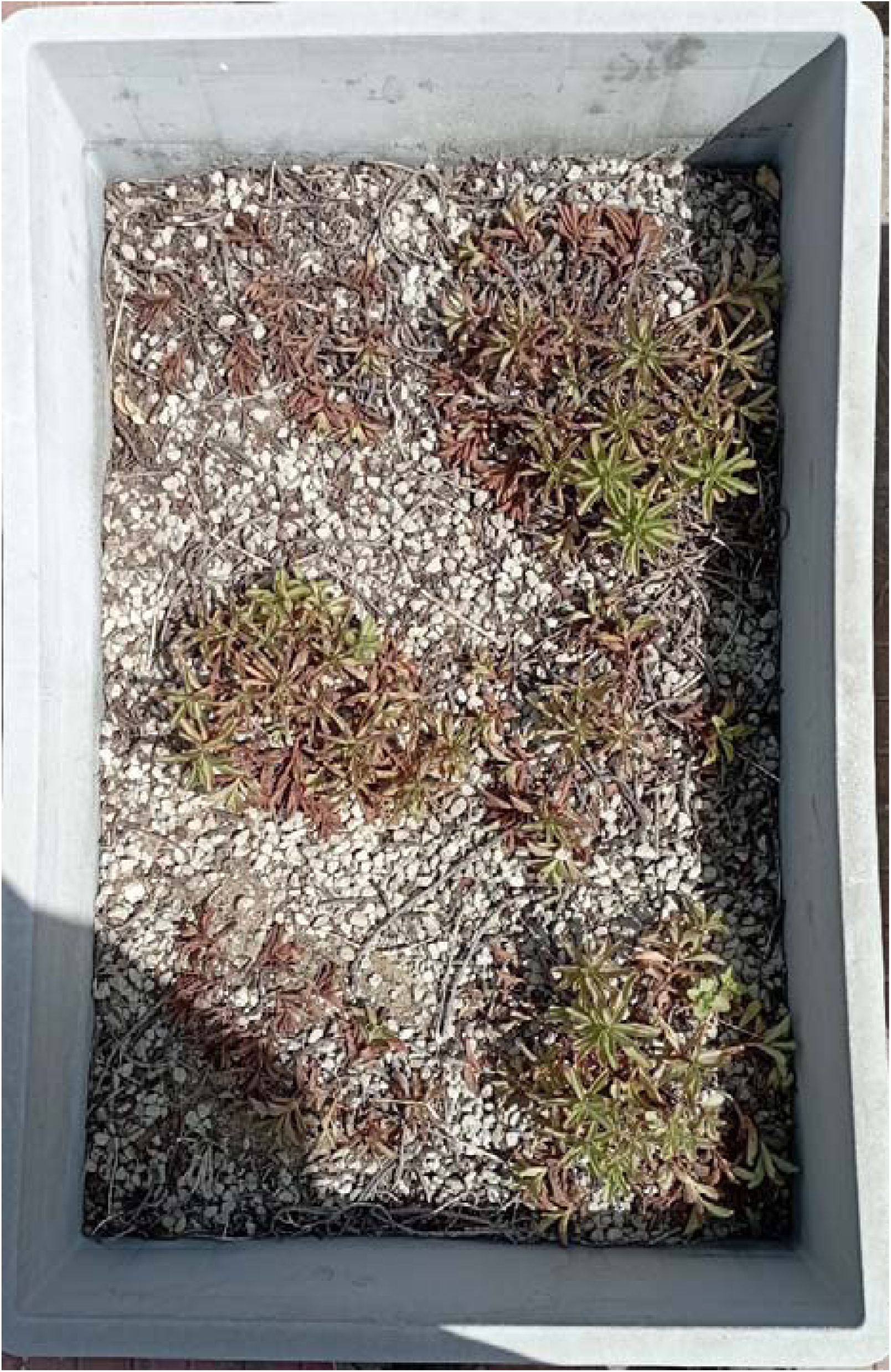
in one of the planting boxes, it is evident that some of the plant communities have shown signs of withering (photographed on July 10, 2023)

Thirdly, it is evident that as a succession study, the time span of 3 years is still too short. Although it can be seen that the succession has gradually stabilized. However, the changes in the PA region are still not significant. If the differences in PA are not significant enough in the next three years, we can have confidence to say that PA is completely unaffected by external succession. It is still too early at present.

## 6. Full text conclusion

6.1 During the epidemic period, the main invasive plants were *Avena fatua Linn. var. fatua, Setaria viridis (L.) P. Beauv, Artemisia capillaris Thunb, Digitaria sanguinalis (L.) Scop*. Except for one planting box with a Shannon index close to 1, most planting boxes only had one invasive plant;

6.2 Natural succession has had a huge impact on the original plants in the rooftop green area, with a significant decrease in average coverage, indicating that the original plants are under pressure from invaders;

6.3 However, the response of the original plants to invasion pressure is not the same, with the SS almost disappearing, while the PA remains lush; And we found that PA seems to have undergone its own succession, that is, its coverage tends to gradually slide towards the average value.

## Acknowledgement

1. Ruoxi Wu, Jingqi Guan, Zihang Xia, Tianqing Zhang: experiment, data collection, paper writing;

2. Yi Jiang, Zhong Wang: literature, data processing, paper examination

3. Thank you for Sirun Chen, Zijian Cui, Jingqi Guan, Yuanze He, Zongyao Shen, Yihe Wang, Yuyuan Wang, Muchen Wei, Zhifan Yang, Guanchen Zhu’s help about experiment of the paper

4. Thank you for Lei Zhao (0000-0003-1643-4981) and Weiqi Zhou (0000-0001-7323-4906)’s help about ideas of the paper

## Notes

### Competing Interest Statement

The authors have declared no competing interest.

